# Place cells in head-fixed mice navigating a floating real-world environment

**DOI:** 10.1101/2020.10.18.344184

**Authors:** Mary Ann Go, Jake Rogers, Giuseppe P. Gava, Catherine Davey, Seigfred Prado, Yu Liu, Simon R. Schultz

## Abstract

The hippocampal place cell system in rodents has provided a major paradigm for the scientific investigation of memory function and dysfunction. Place cells have been observed in area CA1 of the hippocampus of both freely moving animals, and of head-fixed animals navigating in virtual reality environments. However, spatial coding in virtual reality preparations has been observed to be impaired. Here we show that the use of a real-world environment system for head-fixed mice, consisting of a track floating on air, provides some advantages over virtual reality systems for the study of spatial memory. We imaged the hippocampus of head-fixed mice injected with the genetically encoded calcium indicator GCaMP6s while they navigated circularly constrained or open environments on the floating platform. We observed consistent place tuning in a substantial fraction of cells with place fields remapping when animals entered a different environment. When animals re-entered the same environment, place fields typically remapped over a time period of multiple days, faster than in freely moving preparations, but comparable with virtual reality. Spatial information rates were within the range observed in freely moving mice. Manifold analysis indicated that spatial information could be extracted from a low-dimensional subspace of the neural population dynamics. This is the first demonstration of place cells in head-fixed mice navigating on an air-lifted real-world platform, validating its use for the study of brain circuits involved in memory and affected by neurodegenerative disorders.

## 1 INTRODUCTION

Place cells in the hippocampus encode spatial information during navigation, by firing selectively when the animal is in a certain part of its environment (O’Keefe and Dostrovsky, 1971; Muller and Kubie, 1989; Taube, 1995). This location-specific firing requires the encoding and recall of spatial memory, and has been suggested to be the “where” component of episodic memory (Ergorul and Eichenbaum, 2004; Leutgeb et al., 2005; O’Keefe, 2007). As such, it provides a powerful model system for investigating the neural circuit mechanisms underlying learning and memory, as well as the neurological basis of neurodegenerative disorders affecting memory such as Alzheimer’s Disease (Cacucci et al., 2008; Mably et al., 2017).

Place cells have been readily recorded using electrophysiological techniques in freely moving mice and rats. However, additional insight into system function can be gained using single- and multiple-photon fluorescence imaging techniques, which enable large populations of genetically labelled neurons to be monitored simultaneously (Peron et al., 2015; Schultz et al., 2016). While calcium fluorescence place fields in the hippocampus have been imaged using single-photon microendoscopy (Ziv et al., 2013), image quality, imageable depth of field, optical sectioning and consequent cell separability is much greater with two-than one-photon microscopy, which is why it has become the gold standard technique for spatially resolved investigation of cortical circuit function. Two-photon microendoscopy is possible, but is much less well developed, and optical access to the brain is still inferior to that possible in head-fixed preparations (Ozbay et al., 2018; Zong et al., 2017). This has spurred the development of solutions for employing rodent spatial navigation behavioural tasks during head fixation.

One increasingly popular approach is the use of Virtual Reality (VR) platforms. In such systems, a mouse is typically head-fixed atop a polystyrene ball which is free to rotate. Optical sensors tracking ball movement are used to update a visual display (Dombeck et al., 2007; Muzzu et al., 2018), allowing either one- or two-dimensional movement. Place cells have been observed in head-fixed mice navigating in one-dimensional (i.e. linear track) virtual environments in such an apparatus (Harvey et al., 2009; Dombeck et al., 2010; Rickgauer et al., 2014). One advantage of these VR systems is that they allow controlled manipulation of the environment in a way not easily possible in the real world. Moreover, because the mouse’s head is fixed, intracellular recording and two-photon imaging are made feasible. However, there are several key disadvantages of VR systems, including the lack of translational vestibular input, and the lack of sensory feedback of modalities that may be more behaviourally salient for rodents than vision (Ravassard et al., 2015). Two-dimensional place tuning has been shown to be profoundly impaired in VR spatial navigation (Aghajan et al., 2015), and in addition, the theta rhythm frequency has been found to be slower in VR environments (Aronov and Tank, 2014). In fact, 2D place tuning has, to date, only been observed in VR systems where the rodent is suspended in a body jacket attached to a commutator allowing it to make head movements and to rotate its body through a full 360° (Aronov and Tank, 2014), or with a complex commutator headplate attachment that allows head movements constrained to horizontal rotations (Chen et al., 2018, 2019). Neither system allows straightforward extension to two-photon imaging or intracellular recording.

Recently, a 2D real-world system in which mice are head-fixed while navigating a track floating on air has been developed (Kislin et al., 2014; Nashaat et al., 2016). The system allows for sensory feedback and head immobility allows for intracellular recording and two-photon imaging. Until now, the presence of place cells in such an environment has not been shown. We show here that mice can navigate the floating track and give the first demonstration that the system allows for place tuning, with spatial information rates broadly comparable to those observed in head-fixed mice.

## 2 MATERIALS AND METHODS

### 2.1 Animals

All experimental procedures were carried out under the Animals (Scientific Procedures) Act 1986 and according to Home Office and institutional guidelines. Subjects were C57BL/6 mice of age 1 to 9 months at the time of viral injection. Behavioural data were collected from 6 mice (5 males, 1 female, median age: 8.0 months) for the circular track and from 5 mice (all females, median age: 1.5 months) for the circular open field. Imaging data were collected from 3 mice (2 males, 1 female, median age: 8.2 months) for the circular track and from 3 mice (all females, median age: 1.7 months) for the open field. Animals were kept on a reverse 12-h light, 12-h dark cycle with lights on at 7pm.

### 2.2 Virus injection and hipppocampal window

Mice were anaesthetised with 1.5-3% isofluorane. Body temperature was monitored with a rectal thermal probe and kept at 37°C using a heating blanket. Analgesia was administered pre-operatively with Carprofen (5 mg/kg) and buprenorphine (0.07 mg/kg). A small (~0.5 mm) craniotomy was made and the virus AAV1.hSyn1.mRuby2.GSG.P2A.GCaMP6s.WPRSE.SV40 (Addgene 50942, titer 1.9 × 10^13^ vg/ml, −50 nL) was injected into the hippocampus (from bregma, in mm: 1.7-1.8 ML, 2.0 AP) 1.5 mm from the dural surface. The virus contains a green genetically encoded calcium indicator protein (GCaMP6s) and a red fluorescent protein (mRuby) for cell body localisation. Two weeks post-injection, a hippocampal window was implanted as described by Dombeck et al. (2010). A circular craniotomy centred on the previously made injection hole was marked using a 3-mm diameter biopsy punch and the cranial bone was removed using a dental drill. The cortex above the injection site was aspirated using a 27 gauge needle connected to a water pump until the fibers of the corpus callosum became visible. A stainless steel cannula (diameter: 3 mm, height: 1.5 mm) with a glass bottom was then pressed down into the tissue and fixed in place using histoacryl glue. The surrounding skull was roughened using a scalpel blade to make crisscross lines before a stainless steel headplate (aperture: 8.5 mm) was attached to the skull, centred on the craniotomy, using histoacryl glue. Exposed skull outside the headplate aperture was covered with dental cement mixed with black powder paint. Mice were given 5-7 days to recover before behavioural training was started.

### 2.3 Behavioural training

Approximately one week after the hippocampal window was implanted, the animals were habituated to the experimenter by handling and were placed under water restriction. Behavioural training started the following day. Animals were trained to move in the dark either along a circular track (outer diameter: 32.5 cm, width: 5 cm) or in a circular open field (diameter: 32.5 cm), both floating on an air table (Mobile HomeCage Large, Neurotar). Infrared (IR) light illuminated the training area and an IR camera was used to monitor the animals. The floating tracks were made of carbon fibre (weight: 100 ± 2.8 g) and had 4-cm high walls lined with visual (phosphorescent tapes, Gebildet E055 and E068) and tactile cues (sandpaper, cardboard, foam, bubble wrap). The phosphorescent tapes emitted light at 500 (blue) and 520 (green) nm and glowed for the duration of the imaging session (≤ 1 hr per track, Supplementary Fig. 1). The floor of the circular open field was also lined with tactile cues (mesh tape). The air table rests under a two-photon resonant scanning microscope (Scientifica Ltd, Uckfield UK, Fig. 1A) and is fitted with a magnet-based position tracking system that acquires data (including, among others, Euclidean and polar mouse coordinates and speed) at 100 Hz. A lick spout was attached to the headplate mount and automated water delivery was controlled using a peristaltic pump (Campden Instruments). Water rewards were accompanied by a beep.

**Figure 1.**
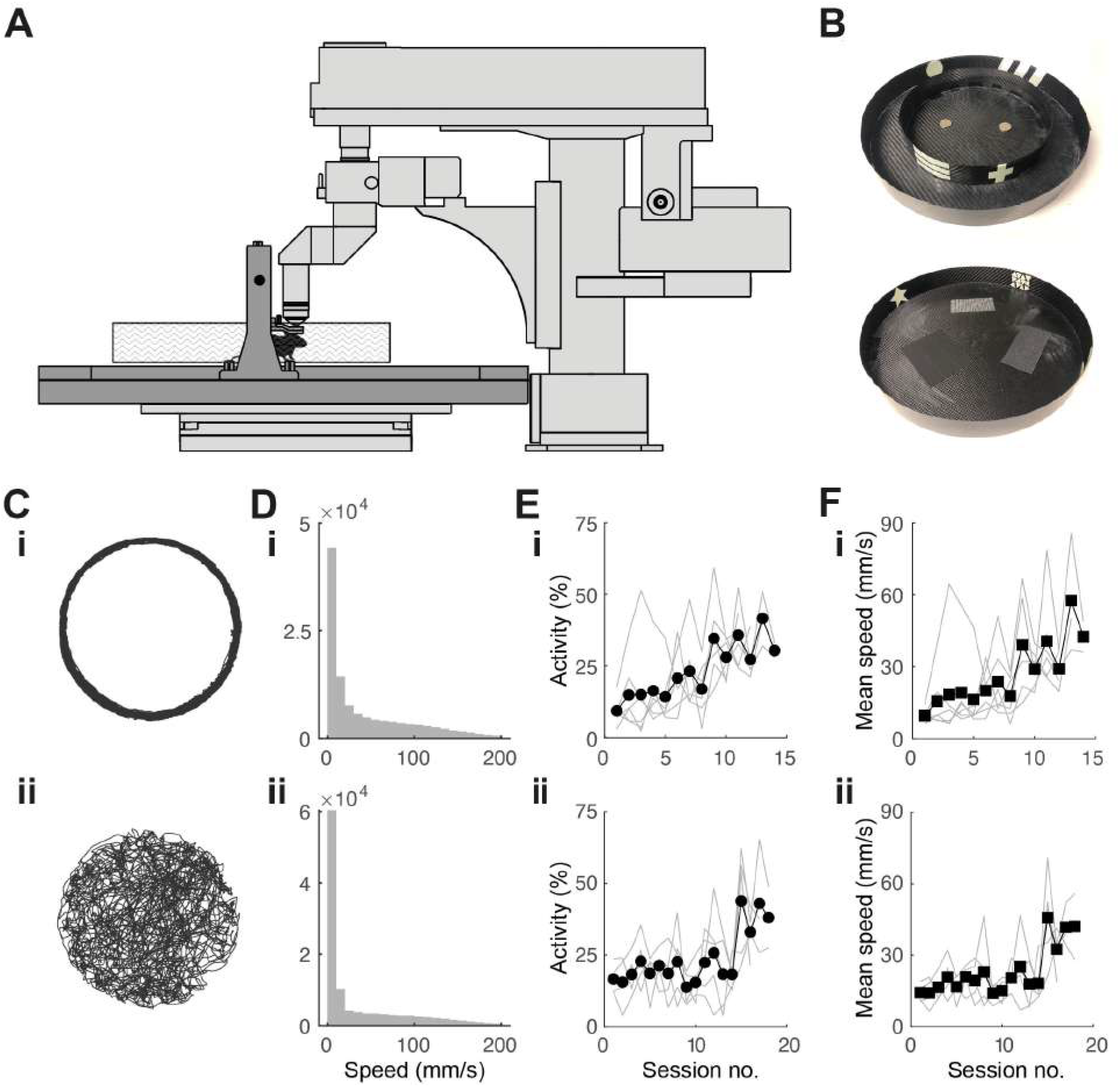
Experimental setup and behavioural training. **(A)** Schematic of experimental setup. Mouse is head fixed while navigating a floating track under a two-photon microscope (in light grey). **(B)** Photographs of circular track (top) and open arena (bottom) used in experiments. Outer diameter for both: 32.5 cm. **(C)** 20-min spatial trajectories of head-fixed mice navigating **(i)** a circular track and **(ii)** a circular open field. **(D)** Distribution of mouse locomotion speeds during the behavioural sessions shown in B. **(E)** Fraction of session spent running in the **(i)** circular track *(N =* 10 mice) and **(ii)** open field *(N =* 5) throughout the training period. **(F)** Progression of running speed in the **(i)** circular track and **(ii)** open field throughout the training period.

Animals were trained twice daily in 45-min sessions. In the circular track, animals were trained with one circular track in the morning and a second circular track with different visual and tactile cues in the afternoon. Animals were given a water reward of 4 uL per loop traversed at random locations. In the circular open field, a 4-uL reward was given every time the animal went from an outer region (outer 6 cm of open field) to either one of two inner regions (two inner 20.5 cm diameter half circles), or vice versa. Daily water intake was limited to 1-3 mL and was individually adjusted for each mouse to maintain the target weight of 85% of the pre-restriction weight. At the start of each training session, two rewards were given to motivate the mice to lick for water. There was no limit to the number of rewards animals could have during sessions. If the animal did not reach the target volume for the day during training, the remaining volume was given at the end of the last training session for the day. Animals were trained for 11-14 sessions in the circular track and 16-18 sessions in the open field before imaging was started. Mice that did not have good GCaMP6s expression in the CA1 region were excluded from imaging experiments.

### 2.4 Two-photon imaging

We used a commercial two-photon resonant scanning microscope (VivoScope, Scientifica) equipped with a tiltable objective mount and a 16× water-immersion objective (LWD 0.8 NA, Nikon). Ultrasound gel at 50% concentration was used as immersion liquid. GCaMP6s and mRuby were excited at 940 nm with a Ti:sapphire laser (Mai Tai, Newport). The laser power underneath the objective was 60-166 mW. Images (512×512 pixels, 330×330 μm or 490×490 μm field of view) were acquired at 30 Hz during imaging sessions which lasted up to an hour. In remapping experiments, mice were imaged for up to an hour in one track then moved to another track where they were imaged for up to another hour. SciScan software (Scientifica) was used for microscope control, and image acquisition was TTL-synchronized to position tracking and reward timing signals. Light from the phosphorescent tapes on the floating track walls used as visual cues was insignificant in the green and red imaging channels.

To characterize stability of place fields, animals were imaged in the circular track in multiple sessions spanning up to 8 days and in the open arena up to 3 days. To locate the same set of cells in later sessions, we used the red channel image from previous sessions as reference.

### 2.5 Processing of calcium imaging movies

We used the MATLAB (Mathworks, Natick, MA) implementation of the CaImAn software package (Giovannucci et al., 2019) for motion correction, automatic identification of regions of interest (ROIs) and deconvolution of neural activity from fluorescence traces. To remove motion artefact from the calcium imaging videos, we first did rigid image registration then non-rigid image registration. For ROI identification, the maximum number of ROIs and the average cell size for a given field of view (FOV) were estimated by examining representative images in ImageJ. Overlapping ROIs were excluded. Signal contribution from the surrounding neuropil was removed using the FISSA toolbox (Keemink et al., 2018). Neural activity was then deconvolved from the neuropil-decontaminated fluorescence traces using the OASIS algorithm (Friedrich et al., 2017). This produced an event train which preserved both the time and amplitude of inferred calcium transient events. In some analyses, the presence/absence of calcium transient events was used; integrating the number of such events over a fixed time window, and dividing by the length of the window yields the neural event rate (units events/sec). In other analyses, we took into account the amplitude of events; in the absence of per-cell calibration to true spike counts, we consider the units of the amplitude of this event train to be (dimensionless) Δ*F/F*, inherited from the original time series extracted from each ROI. Integrating the amplitude of such events over a fixed time window and dividing by the length of the window yields the neural activity rate (units Δ*F/F.s*^−1^).

To track cells across multiple imaging sessions (see Supplementary Fig. 2), we motion-corrected images from different sessions using the motion-corrected image from one session as a template. We then temporally concatenated the videos from different sessions and ran the ROI segmentation algorithm on the concatenated video. For calcium images in the open arena, ROIs across imaging sessions were registered using the CaImAn ROI registration algorithm. ROIs were shifted by registering the templates from individual imaging sessions.

### 2.6 Data analysis

Inferred neural activity and mouse tracking data were analysed using custom scripts written in MATLAB. For place field analysis, only time points for which the animal speed exceeded 20 mm/s were included, to eliminate periods when the animals paused for rewards or for grooming.

For the circular track, mouse position was linearised by converting angular distance to Euclidean distance using the known circumference of the circular track. We used 2-cm bins and computed neural event rates by dividing the total number of neural events during occupancy of the bin by the total occupancy time there. These rate maps were smoothed using a boxcar average over three bins and each map was normalised by its maximum value. Spatial information (in bits/event) for each cell was computed as described previously (Skaggs et al., 1992). Neurons were classified as place cells if they met the following criteria: 1) neural events were present for at least half of the traversals (laps) through the circular track, 2) neural events were present for at least 5% of the time the mouse spent within the place field and 3) the cell contained spatial information greater than chance. Chance-level spatial information for a cell was determined by performing 1,000 shuffles of the time stamps of the inferred neural activity and calculating the spatial information rate for each shuffled neural activity. The cell was considered a place cell if its information rate exceeded 99% of the values for the shuffled data. The location of the place field was defined by the bin location of maximum neural event rate while the place field size was determined by the number of bins for which the neural event rate was at least 50% of the maximum. We quantified stability of place fields across days by calculating recurrence, field correlation and place field shift for all possible pairs of sessions separated by *N* days. Recurrence is the fraction of place cells in one day that retain the classification after *N* days. Place field correlation is the mean correlation of the rate maps of place cells that retained their classification in two sessions N days apart. Place field shift is the difference in field location of place cells in two sessions at N days interval.

For the circular open field, bins were 2 cm × 2 cm and the rate maps were smoothed with a 2D Gaussian smoothing kernel with a standard deviation of 1.5. Neurons were classified as place cells if the cells contained spatial information greater than chance. To eliminate single bursts of activity generating place cells, neural events had to be present for at least 2% of the time the mouse spent within the place field. Place field location was obtained by calculating the centroid location of the normalised rate map above a threshold of 0.5.

In the Results that follow, we describe estimates of the means of distributions by the sample mean ± the standard error of the mean, unless otherwise indicated. One exception to this is in the reporting of mean activity rates (Δ*F/F.s*^−1^), which are expected to fall (approximately) on a log-normal distribution, thus a symmetric distribution of errors would be inappropriate even in the limit of large samples. In this case, therefore, we instead report the 90% confidence interval of the mean.

Neuronal variability analyses were performed by adapting methods from classical visual neuroscience literature (Tolhurst et al., 1983) to calcium transient amplitude-event trains; instead of measuring how variable neuronal responses were across repeated presentations of a stimulus, we analyzed how variable responses were to laps around the same track. For each cell, we accumulated the amplitude of calcium transient events occurring when the mouse was in each 2-cm bin, dividing it by the amount of time spent in that spatial bin. Averaging or taking the variance of this quantity across laps gives the mean activity or variance, respectively, for that spatial bin; as the mouse progresses along the track (e.g. into and out of a place field), the mean changes, and thus we obtain the relationship between variance and mean of the activity for that cell.

We used total least squares linear regression to fit a power law model *y = ax^β^* to the relationship between activity variance and mean for each individual cell. The power law exponent β, which can be read off from the slope of the fitted line, provides useful information about the reliability of neuronal signalling. An exponent of 1 indicates reliability equivalent to that of a Poisson process; above 1 implies additional sources of variability.

To study neural population dynamics, we embedded the population calcium traces, recorded during periods when mouse running speed exceeded 20 mm/s, in a manifold to extract lower-dimensional population activity patterns. Given the population activity matrix **X** of size *N × T*, where *N* is the number of recorded units and *T* the number of time samples, there are *T* population vectors **x**_*t*_ of length *N*. The aim of this analysis was to uncover a lower-dimensional embedding **Y** (*M × T*), where *M* ≪ *N* is the number of manifold dimensions. To do this, we used Classical Multidimensional Scaling (MDS), after observing qualitatively and quantitatively (in terms of variance accounted for) better representation (Supplementary Fig. 3) in comparison to Principal Components Analysis (PCA). We generated a dissimilarity matrix to account for the difference between population activity vectors **x**_*t*_ over the entire time course *T*. We used the cosine distance metric, which evaluates the angle between any two population activity vectors **x**_*t*_ at any two time points. ISOMAP produced a similar embedding (Supplementary Fig. 3). For the purpose of comparing manifold dimensionality, we defined the dimensionality as the number of manifold components (MDS eigenvalues and associated eigenvectors) required to explain 90% of the variance in the population activity.

To decode the mouse’s angular position (in either the circular track or open field environment), we applied an Optimal Linear Estimator (Muzzu et al., 2018), incorporating a varying number of manifold dimensions, and using 5-fold cross-validation.

## 3 RESULTS

### 3.1 Mouse behaviour in floating environment resembles tethered and free behaviour

Head-fixed mice were trained to navigate the floating track system (Fig. 1A) in the dark using operant conditioning. We designed visual cues on the environment walls using phosphorescent tapes. These were visible in the dark for the duration of the training session (45 min) and did not add significant weight to the tracks, which would have increased their rotational inertia and made them harder to control. The outer diameter (32.5 cm) of the floating track was constrained by the distance from the objective lens to the back wall of the microscope.

In the circular track, the mice initially displayed minimal activity. They struggled to propel the track with their feet, tracing circular trajectories that were irregular. Without any intervention, the mice quickly adapted to moving in smooth circular trajectories (Fig. 1Ci) running up to speeds of 200 mm/s (Fig. 1Di). Activity and average speed increased after about 9 sessions (Fig. 1Ei, Fi). By the end of the training period, mice could complete >100 laps in 45 min.

In the circular open field, our goal was for the trajectory of a mouse to completely cover the 2D environment, an important requirement for place field analysis. We rewarded the mice when they went from an outer region (outer 6 cm of open field) to either one of two inner regions (inner 20.5 cm diameter half circles) and vice-versa and found this protocol to be effective in achieving the desired behaviour. The mice adapted to controlling the circular track with their feet, tracing meandering trajectories that were sometimes near the wall and sometimes crossed the arena. This behaviour resembles those of freely-moving (Benjamini et al., 2011; Samson et al., 2015) and tethered (Dupret et al., 2010; Trouche et al., 2016) mice exploring circular arenas (Fig. 1Bii). Mice sometimes showed preference for peripheral locations in the open field. Locomotion speeds were similar to the circular track (Fig. 1Cii), although it took longer for the mice to increase activity and average speed, requiring 15 or so sessions (Fig. 1Dii, Eii). To quantify coverage, we divided the circular arena into 2 cm × 2 cm bins. By the end of the training period, mouse trajectories covered >90% of the open field in 45 min.

Mice were trained in the floating track system after viral injection and hippocampal window implantation. The similarity of their behaviour to that’s observed in tethered and freely-moving mice is validation that our surgical procedures do not adversely affect behaviour, consistent with previous reports (Dombeck et al., 2010; Pilz et al., 2016). Greater numbers of animals were available at earlier than later stages of the experimental pipeline simply because of the requirement to pass successive criteria relating to behavioural performance, quality of preparation, GCaMP6s expression located in area of interest, and registration of ROIs over multiple sessions.

### 3.2 CA1 cells form reliable 1D place fields in a floating circular track

To optically record the activity of CA1 neurons, we injected the adeno-associated virus AAV1.hSyn1.mRuby2.GSG.P2A.GCaMP6s.WPRSE.SV40 into the hippocampus, removed the overlying cortex and implanted an imaging window. Two-photon imaging (at 940 nm) through the hippocampal window three weeks later showed robust expression of both the calcium indicator GCaMP6s and the static marker mRuby in a large population of CA1 neurons (Fig. 2A). We primarily used the mRuby image to repeatedly identify the same cells across time.

**Figure 2.**
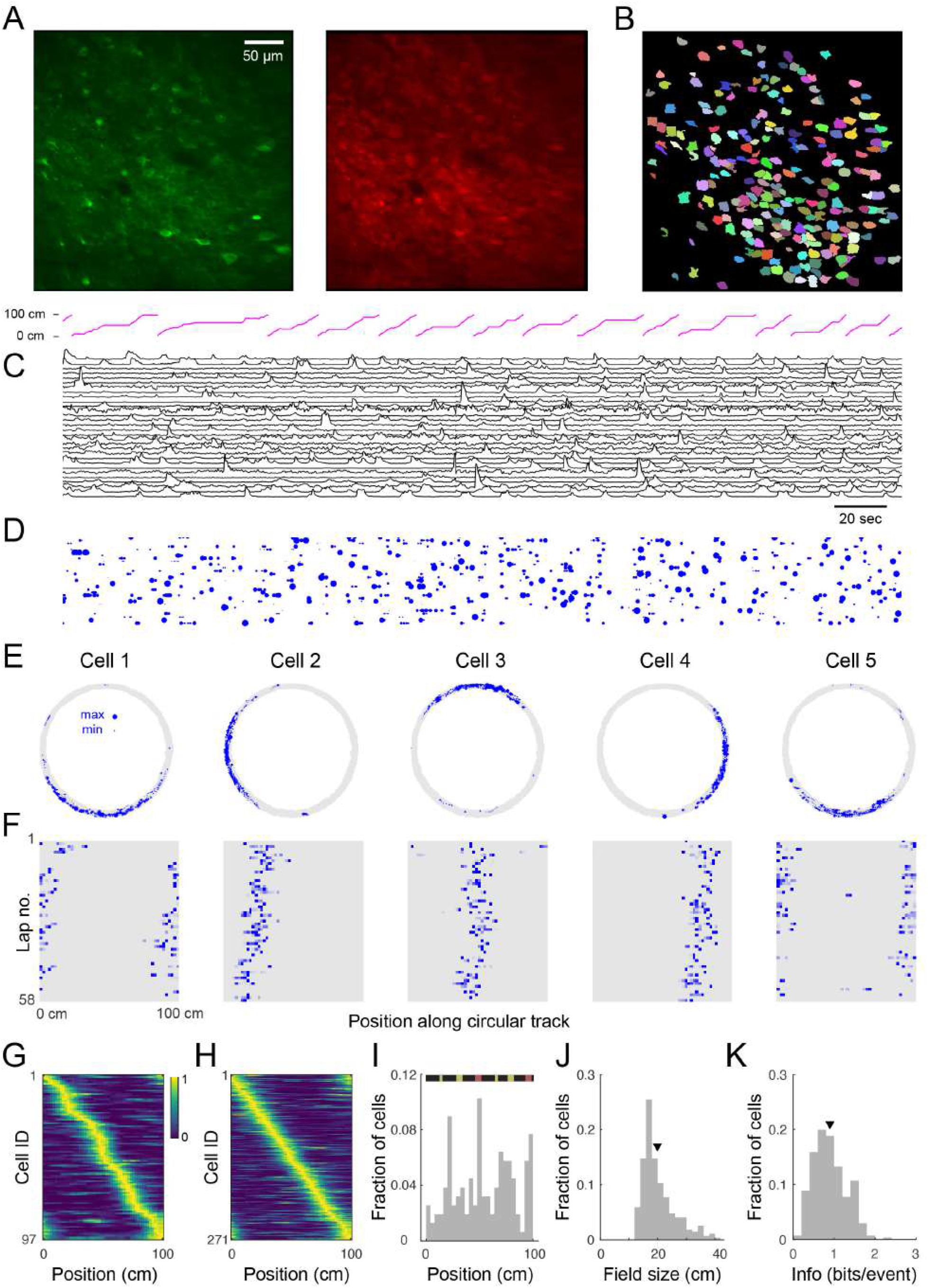
Place cell firing during navigation of a circular track. **(A)** Two-photon images from a typical CA1 imaging session. Green: GCaMP6s, red: mRuby. **(B)** 225 segmented ROIs from image in **A**. **(C)** Ca^2+^ transients for 30 (of 225) randomly selected cells from the region shown in **A**. Trace height is normalised to the 99th percentile of the Δ*F/F* for each cell. Magenta trace at top shows position of the mouse along the circular track. **(D)** Rastergram showing events detected from **C**. Blue dots indicate the time of onset of each Ca^2+^ transient, with dot area showing relative event amplitude (normalised for each cell). **(E)** Spatial trajectory over 24 minutes of recording (gray line) together with locations of firing (blue dots) for five representative cells from the region shown in **A**. **(F)** Neuronal activity maps for the cells in **E** across repeated laps, obtained by integrating the inferred activity in each spatial bin, and normalising by the maximum for each cell. **(G)** Normalised neural event rate maps for 97 (of 225) cells from the region shown in **A** considered place-sensitive, sorted by place preference (peak location). **(H)** Normalized place field map for 271 place cells pooled from 6 fields of view (FOVs) from 3 mice, sorted by place preference. **(I)** Distribution of place field locations in a familiar track. (Top) Schematic of linearised circular track. Visual cues appear in yellow and magnetic doors in red. The inner door is in the middle and the outer door is at the end. (J-K) Distributions over all cells in **H** of place field size **(J)** and spatial information (calculated via Skaggs approximation) **(K)**. Triangles in this and later plots denote mean values. Mean place field size (± s.e.m.): 19.7 ± 0.4 cm, mean spatial information: 0.90 ± 0.02 bits/event.

To verify place coding in CA1 neurons, we acquired two-photon time-series videos of GCaMP6s fluorescence in the hippocampi of mice running along a circular track. We used an automated algorithm (Giovannucci et al., 2019) that identifies cells and extracts their activity from the GCaMP6s fluorescence changes (Δ*F/F*). We found 53-225 (median: 101) active neurons per FOV (330×330 μm, 6 FOVs in 3 mice) imaged for 12-28 min (see Fig. 2B showing 225 ROIs for the representative image shown in Fig. 2A). We extracted Δ*F/F* traces from the ROIs and observed significant calcium transients (Fig. 2C). We then used the deconvolved neuronal activity as a measure of spiking activity (Fig. 2D).

In total, we analysed 721 cells from 6 imaging areas in 3 mice and found that 12-43% (median: 35%) of the detected neurons in each imaging area showed location-specific activity characteristic of place cells (Fig. 2E). These cells had well-defined fields of neuronal activity which were apparent with repeated traversals (laps) of the circular track, though not occurring in every lap (Fig. 2F). Moreover, similar to reports in rats (Mehta et al., 1997; Lee and Knierim, 2007), we observed in some cells a backward shift in the location of the place field in later laps (Fig. 2F). Place cells were active in 72.1 ± 0.6% (mean ± s.e.m.) of the laps and their firing fields covered the length of the circular track, although non-uniformly distributed across it (Fig. 2G-I). Many of the place fields mapped positions preceding or within local cues (Fig. 2I) as previously reported (Geiller et al., 2016). Place cells also showed a skewed distribution of place field size with an average of 19.7 ± 0.4 cm (Fig. 2J). On average, the information rate of place cells was 0.90 ± 0.02 bits/event (Fig. 2K), within the range of values reported for freely moving mice (Chen et al., 2013; Mou et al., 2018; Gonzalez et al., 2019) and for head-fixed mice navigating virtual linear tracks (Arriaga and Han, 2017).

Imaged hippocampal CA1 neurons produced an average of 0.65 ± 0.01 calcium transient events per second *(n* = 721 cells) during locomotion around the circular track. This is consistent with previous observations from freely-moving mice (McHugh et al., 1996), given that many of the calcium transients we measure likely result from calcium influx due to multiple action potentials. Place cells fired at higher average rates than non-place-sensitive cells (mean 0.95 ± 0.02 events/sec, *n* = 271 versus 0.46 ± 0.01 events/sec, *n* = 450, respectively; significance level 2 × 10^−81^, one-sided Student’s t-test). The distribution of place cell mean neural event rates during the session was skewed towards higher rates (Fig. 3A). The distribution of mean neural activity rates, however (i.e. the rate of events weighted by the amplitude of each calcium transient), was apparently more close to being log-normally distributed across cells (Fig. 3A), with a skew towards lower activity contributed by lower activity non-place cells, consistent with data from extracellular recordings (Buzsaki and Mizuseki, 2014). While a Kolmogorov-Smirnov test of the overall distribution of activity rejected the null hypothesis of log-normal distribution due to this low tail *(p* = 7 × 10^−7^, *n* = 721), the log-normal model could not be rejected for the distribution of place cell activities (*p* = 0.7, *n* = 271).

**Figure 3.**
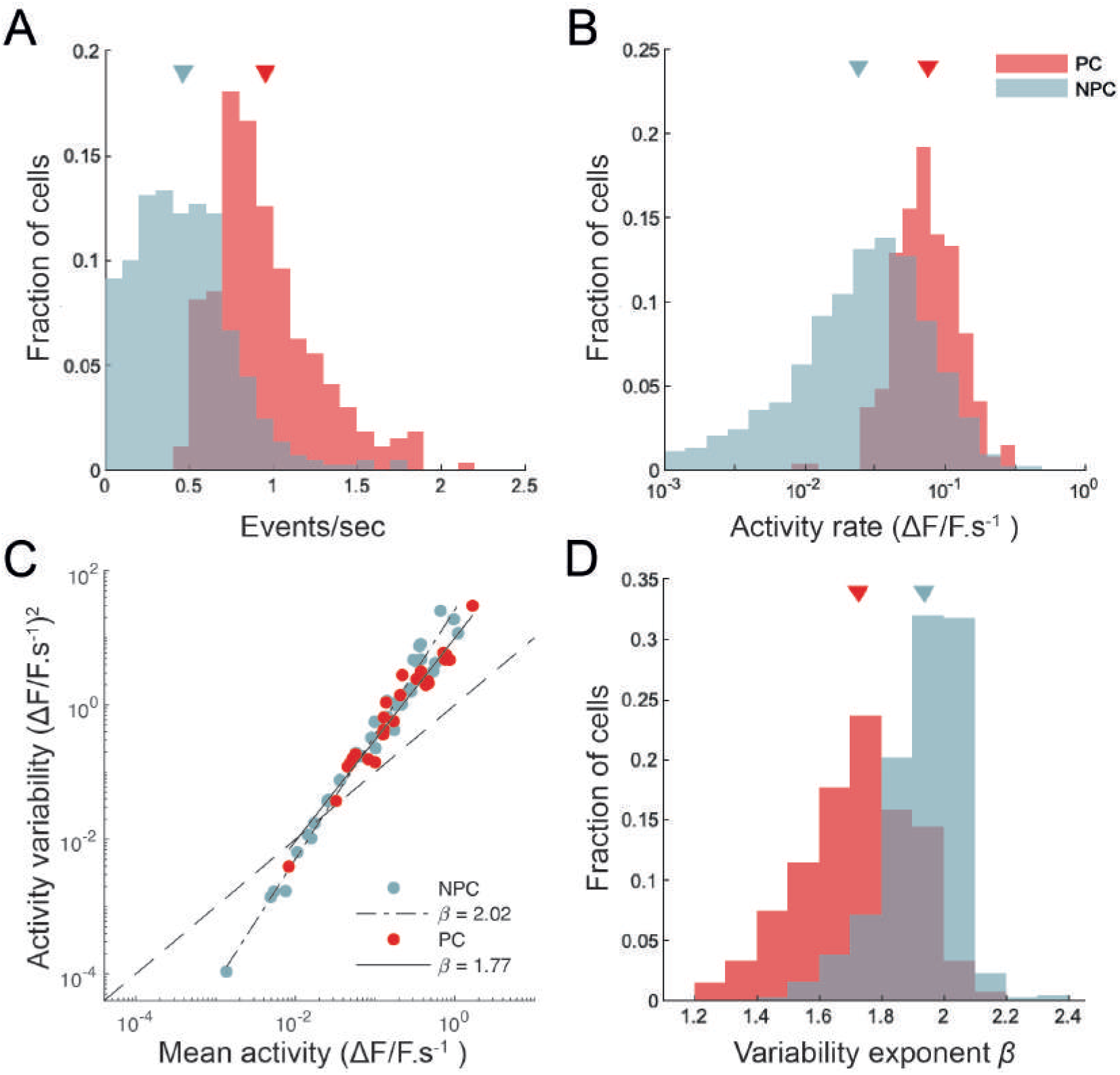
CA1 place cells show higher and more reliable activity than non-place cells. **(A)** Histograms of calcium transient detected event rates for place cells (PC) and non-place-sensitive cells (NPC). Sample means for each group indicated by triangles. **(B)** Histograms of activity rates for place and non-place cells. Activity rates are calculated here by integrating all calcium transient events across a lap, weighted by the amplitude of each event, and dividing by the time taken for the mouse to complete the lap. **(C)** The variance of the activity rate (per unit time spent) in each 2-cm spatial bin as the mouse completed the lap plotted against the mean activity rate, for two cells, one place-sensitive and other not. Total least-squares regression fits are shown. This plot is double-logarithmic, and the slope of the linear fit thus indicates the exponent in the relationship between the variability and mean. **(D)** The distribution of variability exponents for place and non-place cells.

The reliability of neuronal activity has often been measured in the sensory neuroscience literature (Tolhurst et al., 1983; van Steveninck et al., 1997) by calculating the ratio of the variance to the mean of the number of spikes fired in response to each identical stimulus presentation. We are not aware of a similar analysis in the hippocampal literature (Markus et al., 1995, performed a related but somewhat different analysis), however the degree of control our head-fixed preparation affords makes such an analysis feasible. We measured the mean and variance over repeated laps of the neural activity rate for each cell, as described in Methods. Fig. 3C shows the neural activity variance to mean relationships for a pair of cells (one a place cell, the other not), together with power law fits. The exponent of the power law fit (slope of the straight line on a double-logarithmic plot) captures the intrinsic variability of the cell’s response. We found that places cells had systematically lower exponents (i.e. more reliable activity from lap to lap, taking into account the mean level of activity) than did non-place cells (Fig. 3D); mean exponent 1.72±0.01, *n* = 269 for place cells, and 1.94 ± 0.01, *n* = 448 for non-place cells). Although place cells were significantly more reliable from trial to trial than non-place cells, our results suggest that CA1 activity, as measured from calcium fluorescence, is somewhat more variable from trial to trial than neocortical activity (Tolhurst et al., 1983).

### 3.3 Place cells remap almost completely between arenas

Place cells have the ability to change their activity patterns in different environments, a phenomenon known as global remapping. To determine whether place tuning in our floating track is environment-dependent, we imaged mice in a circular track with visual cues (track A) for up to 24 min then transferred them to a second circular track with distinct visual cues and additional tactile cues (track B) where they were imaged for up to 28 more minutes. We found that upon switching to track B, the location-specific activity of many cells disappeared, indicating remapping (Fig. 4A-C). We analysed 154 cells (from 3 imaging areas in 3 mice) which were active in both tracks. 58 of the cells had place fields in both tracks, only 3% of which retained their place field location upon switching to track B. The neuronal activity patterns in the second track could not be predicted from their neuronal activity in the first track – mean activity correlation between tracks was 0.02 ± 0.02. Moreover, place field locations in track B appeared to be randomly redistributed around the circular track relative to their positions in track A (failure to reject the null hypothesis of a circular uniform distribution of place field shifts, *p* = 0.95, Hodges-Ajne test, *n* = 103; Fig. 4C). Altogether, these results show global remapping as a result of visual and tactile cues providing sufficient sensory context for mice to identify the tracks as being different environments despite their identical geometry and dimensions. Similar remapping of place fields in different environments has been previously reported in freely-moving rodents. (Muller and Kubie, 1987; Anderson and Jeffery, 2003; Arriaga and Han, 2017).

**Figure 4.**
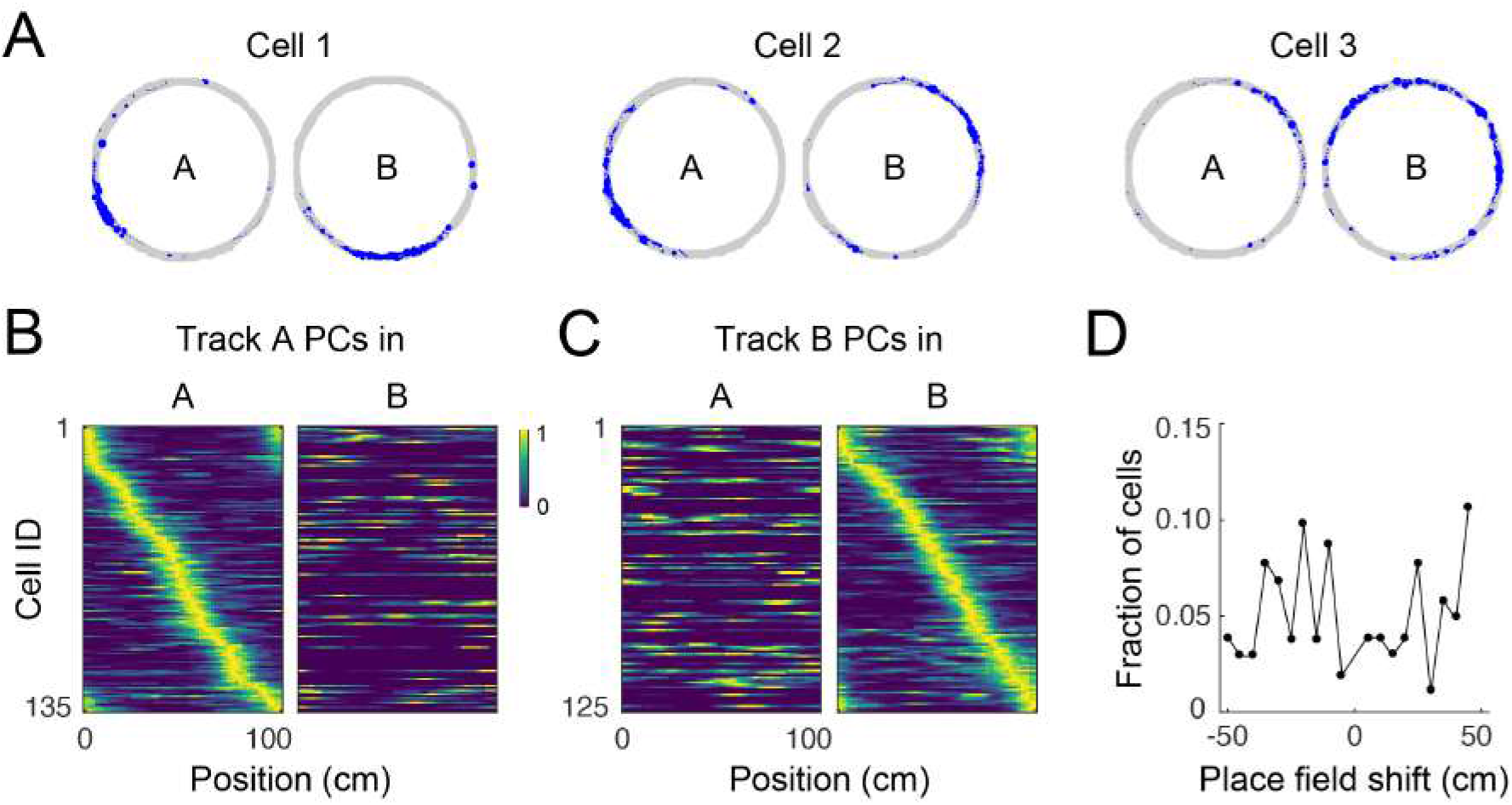
Place cell remapping. **(A)** Spatial trajectory over 24 minutes of recording (gray line) and firing locations (blue dots) for three representative cells in a mouse that navigated two circular tracks with distinct visual and tactile cues. Note that spatial activity of cells in track A is not maintained in track B, indicating remapping. **(B)** Normalised neural event rate maps for track A place cells (PCs) during navigation in track A (left) and B (right) sorted according to place field locations in track A. Cells pooled from three imaging areas in three mice. **(C)** Normalised neural event rate maps for track B PCs during navigation in track A (left) and B (right) sorted according to place field locations in track B. **(D)** Shift in place field locations for place cells common to both tracks.

### 3.4 Dynamic place fields over days

To study the stability of place fields over days, we imaged mice in the circular track multiple times over 8 days. Place cells had consistent location-specific activity within a session but the location was not constant over days (Fig. 5A). For each mouse, there was a similar number of place cells across sessions and the set of place fields for each session fully covered the length of the circular track (Fig. 5B). We found a total of 803 cells in the FOV across all imaging sessions, of which 88 ± 2% were active on any day (Fig. 5C), within the range of reported values for freely-moving mice (Gonzalez et al., 2019). On average, 29 ± 3% of the active cells in each session were place sensitive. Only 4 ± 1% of all cells were place sensitive in all sessions. If a cell was active on one day, the probability that the cell would be active in a later session (here called recurrence) did not change with time (Fig. 5D) within the time period that we examined (< 8 days). Similarly, if a cell was place-sensitive on one day, the probability of the cell being place-sensitive in a later session did not change with time (Fig. 5D). Imaging over a much longer timescale (30 days), Ziv et al. (2013) observed a decrease in the recurrence probability of active and place cells with time. Of the place cells that retained the classification at a later session, there was substantial remapping (Fig. 5E). The decline in similarity of place fields (i.e. substantial remapping) across our imaging period (Fig. 5B,D) is comparable to the observed decline in VR studies imaged within a similar time period (Hainmueller and Bartos, 2018) and is more pronounced than the observed decline in freely-moving mice (Ziv et al., 2013; Gonzalez et al., 2019) imaged over a longer time period (10-30 days). Remapping of place fields over time was consistent with a decrease in place field correlations (mean correlation of the rate maps of place cells that retained their classification in two sessions N days apart, Fig. 5F), similar to reports for head-fixed mice in VR (Hainmueller and Bartos, 2018).

**Figure 5.**
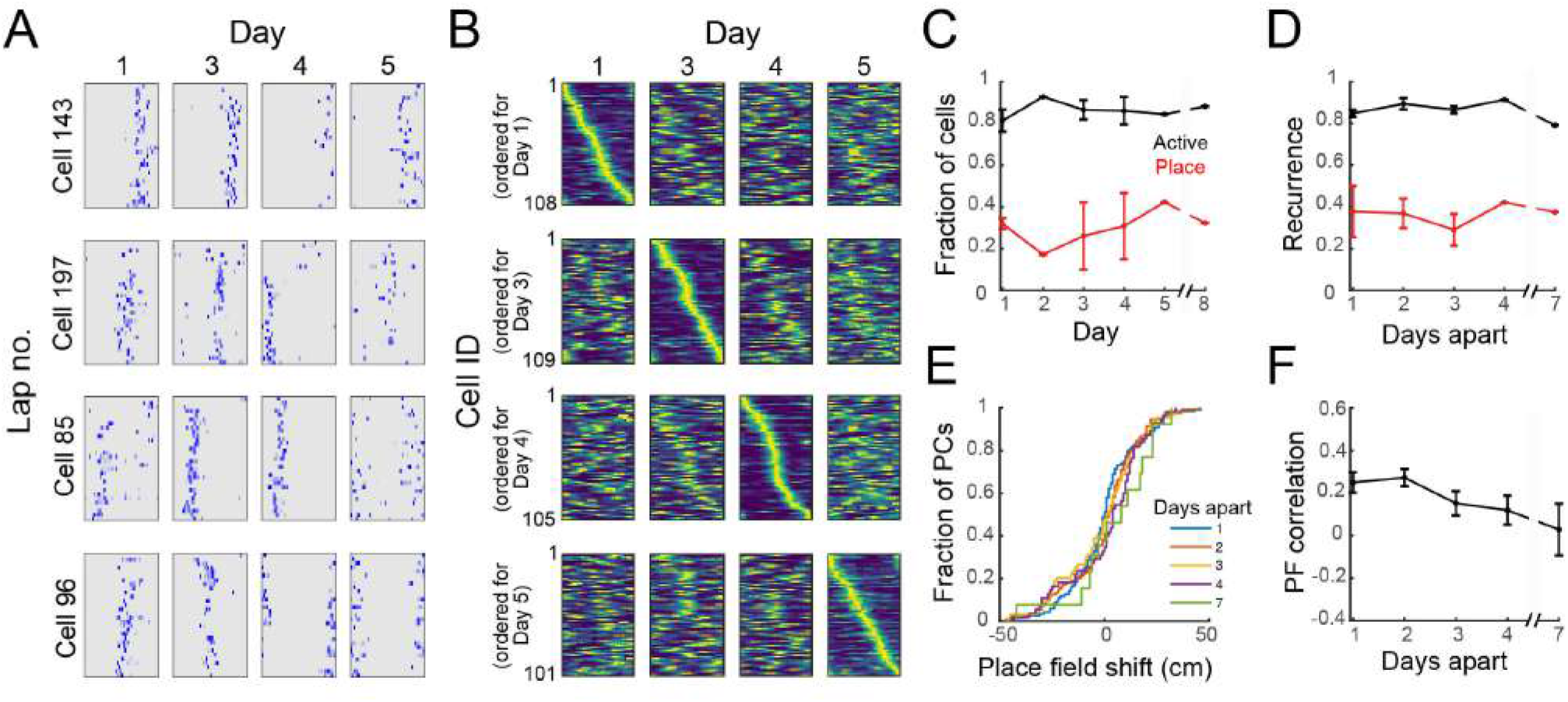
Dynamic place fields. **(A)** Neuronal activity maps across laps in a session and across imaging days for 4 representative cells. **(B)** Normalised place field maps for place cells found on multiple days ordered by maximum location on different days; data from one representative mouse. **(C)** Proportion of all cells found in all imaging sessions that were active on each day and proportion of active cells on that day that were found to be place-sensitive. **(D)** Fraction of cells that were active or place-sensitive in one session which retained that classification at a later session (recurrence) versus number of days interval between the imaging sessions. **(E)** Cumulative distribution of place field shifts for 1 to 7 day intervals between imaging sessions. **(F)** Place field correlation for cells that retained their classification in two sessions versus number of days interval between the imaging sessions. (C-E) Data pooled from 4 imaging areas in 3 mice. Where an error bar is missing, the average was taken from a single imaged area.

### 3.5 Place fields emerge in a floating open arena

In view of reports that spatial selectivity is impaired in two-dimensional virtual environments (Aghajan et al., 2015), we were interested to see whether 2D place fields could be observed in the floating track system. To investigate this, we imaged mice navigating an open circular arena for 20-36 min. We found 44-301 (median: 231) active neurons per imaged FOV (490 × 490 μm, 4 FOVs in 3 mice, 807 cells in total). 16-46% (median: 25%) of these cells showed location-specific activity characteristic of place cells (Fig. 6A)). These activity patterns and place field maps are qualitatively similar to real-world systems (Cacucci et al., 2007; Renaudineau et al., 2009; Ziv et al., 2013) and VR systems that allow rotations of the head or body (Aronov and Tank, 2014; Chen et al., 2018). Some cells responded to the presence of the wall of the arena where landmarks were located, firing near them, resembling border cells that have been observed in other regions of the hippocampal formation in rats (Solstad et al., 2008; Lever et al., 2009; Boccara et al., 2010). Place field centroids were non-uniformly distributed over the arena. Many place fields mapped locations to the side of visual cues and close to tactile cues (Fig. 6B). The bin with the most place fields was at the center of the arena, surrounded by three tactile cues. Overall, mice spent 59 ± 8% of the time in the outer 6 cm of the arena and a greater proportion of cells had place fields with centroids located in this outer region than in the inner region (59% vs 41%). Place cells had a skewed distribution of field size with an average of 172 ± 6 cm^2^ or 21 ± 1% of the environment (Fig. 6D). The distribution of spatial information was similarly skewed with an average of 0.94 ± 0.03 bits/event (Fig. 6E). This value is within the range reported for freely-moving mice navigating real-world systems (Cacucci et al., 2008; Renaudineau et al., 2009; Rochefort et al., 2011) and is greater than that reported for head-fixed mice navigating VR systems with horizontal head rotation (Chen et al., 2018).

**Figure 6.**
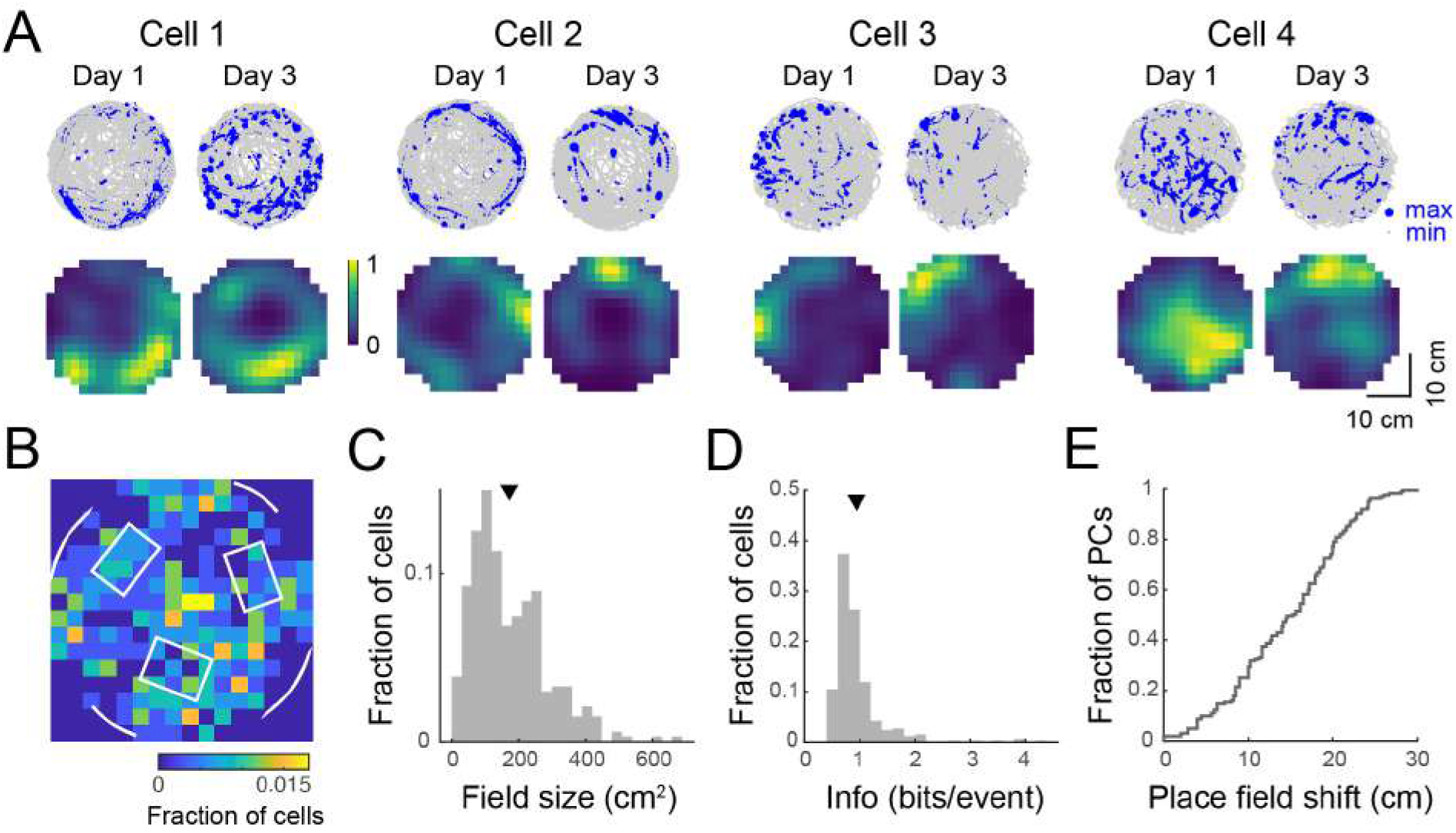
Place tuning for open field behaviour. (**A**) (Top) Spatial trajectory over 20-32 minutes of recording (gray line) together with firing locations (blue dots) for 4 representative cells from 2 mice imaged across two sessions. (Bottom) Normalised place field maps for the cells shown above. (**B-D**) Distributions over 334 cells (pooled from 4 imaging areas in 3 mice) of place field centroid location (**B**, white overlay: schematic of visual cues on the wall and tactile cues on the floor), place field size (**C**), and spatial information (**D**). Mean place field size (± s.e.m.): 172 ± 6 cm^2^, mean spatial information: 0.94 ± 0.03 bits/event. (**E**) Cumulative distribution of place field shifts between the two imaging sessions.

When we imaged mice over two sessions two days apart, we observed the same substantial remapping (Fig. 6A,E) that was evident in the linear track over a period of several days. For recurring place cells, the mean correlation of their place field maps over the two sessions was 0.28 ± 0.04.

### 3.6 Spatial location is accurately encoded in low dimensions of the neural manifold

To study how spatial behaviour on the air-lifted platform is reflected in neural population dynamics, we performed a neural manifold analysis using the MDS technique. Applying this to data recorded from mice running in the circular track revealed distinct recapitulation of the spatial trajectory in the low order components of the neural manifold (typical example shown in Fig. 7A). This result was not specific to MDS, and very similar results were observed using PCA and Isomap (Supplementary Fig. 3). Open field exploration, being less constrained, unsurprisingly revealed much more complex dynamics on the manifold. Colour-coding points (corresponding to patterns of activity across the observed neurons) by angular location around the circular track revealed some clustering of points in manifold localities, spread however throughout the manifold (Fig. 7B; typical example). Colour-coding instead for radial position revealed additional structure at larger scales (Fig. 7C). Variance in the neural population dynamics was accounted for by a smaller number of components in circular track than open-field experiments (Fig. 7D); or equivalently, to account for a fixed amoount of variance required more components (dimensions) for mice exploring an open-field. Considering the number of components required to account for a fixed percentage (90%) of the variance as the ‘‘dimensionality” of the manifold ((Stringer et al., 2019)), we examined how the dimensionality grew with the number of neurons incorporated within an ensemble. This grew at a faster rate with ensemble size for the circular track than open-field environments (Fig. 7E), suggesting that the greater complexity of the exploration trajectory was recapitulated in higher-dimensional neural population dynamics. Finally, we assessed the extent to which spatial information could be decoded from low-dimensional neural manifolds, by training an Optimal Linear Estimator (OLE) decoding algorithm on the angular position data. Angular position could be decoded with high accuracy from a very small number of components in the circular track case (Fig. 7F); in comparison, decoding of angular position in the open-field task yielded lower fidelity estimates, and required more manifold dimensions.

**Figure 7.**
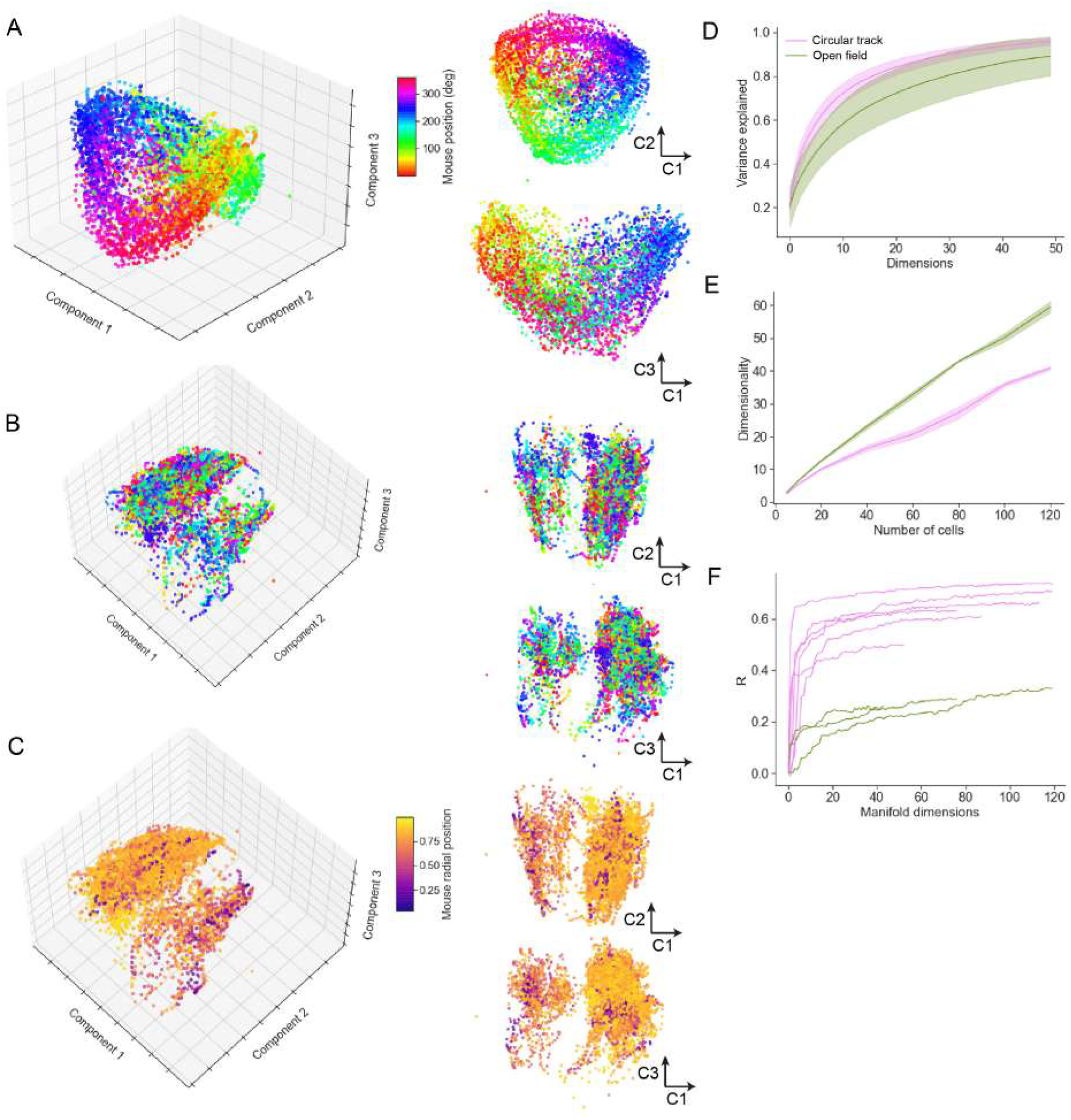
Neural manifold analysis reveals mouse spatial trajectories in a low-dimensional subspace of the neural population dynamics. (**A**) Multidimensional Scaling (MDS) analysis of a typical circular track experiment. The first three components shown in three-dimensional projection at left, with 2D projections at right. Each dot reflects the projection of the neural population activity at a given time point; colour code indicates spatial location in degrees of angle around the track. (**B**) MDS depiction of a typical open field experiment. In this case the colour code is as in (A), reflecting angular but not radial location. (**C**) Remapping the colour code to reflect normalised radial location indicates additional manifold structure relating to spatial position. (**D**) The variance explained by inclusion of an increasing number of manifold dimensions for the circular track and open field environments. (**E**) The growth of dimensionality, defined as the number of components needed to capture 90% of the variance, with the number of cells included in the manifold. (**F**) Decoding performance, measured by Pearson correlation coefficient, for circular track and open field environments.

## 4 DISCUSSION

We have here provided the first demonstration of hippocampal place tuning in head-fixed mice navigating a real-world floating track. Our results indicate reliable place fields in a one-dimensional circular track, comparable to those observed in freely moving (McHugh et al., 1996; Chen et al., 2013) and head-fixed mice navigating a virtual reality apparatus (Dombeck et al., 2010), with spatial information rates similar to both (Arriaga and Han, 2017; Mou et al., 2018; Gonzalez et al., 2019). When mice re-entered the same environment, place fields were recapitulated, with fields dynamically changing over the course of days, with a rate of reconfiguration similar to that observed in VR recordings (Hainmueller and Bartos, 2018), but significantly faster than that seen in real-world free exploration (Ziv et al., 2013; Gonzalez et al., 2019).

We also demonstrated the presence of 2D place fields in an open arena, similar to those seen in mice during free exploration (McHugh et al., 1996; Cacucci et al., 2007; Renaudineau et al., 2009; Ziv et al., 2013) and VR systems that allow rotations of the head or body (Aronov and Tank, 2014; Chen et al., 2018), with spatial information rates similar to the former. We initially found this surprising, as the lack of vestibular sensory input might be expected to impair place cell formation (Stackman et al., 2002; Aghajan et al., 2015). However, the abundance of multimodal sensory information in the floating track overcomes the lack of vestibular signals. These results strengthen the hypothesis that rather than there being a strict requirement for vestibular cues, the repeated pairing of multisensory inputs is key to strengthening spatial selectivity (Aghajan et al., 2015). We speculate that the lack of vestibular information may likewise affect the properties of the motion-dependent theta rhythm in the floating track system, similar to observations from VR (Aronov and Tank, 2014; Ravassard et al., 2015; Chen et al., 2018), although this has yet to be verified. Many hippocampal researchers are skeptical of the utility of VR for studying spatial navigation, with Donato and Moser (2016) commenting on the Aghajan et al. (2015) results: “These data cast doubt on whether the way animals interpret 2D or 3D space can ever be understood using VR under conditions of head or body restriction. Strategies that compensate for the loss of synchrony between vestibular information and the animal’s behaviour would be a welcome advance”. Our results here suggest that the availability of a richer, multimodal sensory scene can go some way towards addressing this issue.

The floating track system has several advantages over VR systems. Notably, it allows the expression of 2D place cells in head-fixed setups without the need for a complex commutator headplate (Chen et al., 2018), making two-photon imaging straightforward. In addition, although not as dynamically reconfigurable as pure VR systems, the floating track system employs multimodal sensory stimuli which contribute to the integrated spatial context (Ravassard et al., 2015; Geva-Sagiv et al., 2015), making it adaptable to a variety of tasks. Multimodal VR systems have been developed which use distal visual and auditory stimuli (Cushman et al., 2013). In contrast, the floating track has the capability to employ local visual, auditory, tactile and olfactory cues. Studies have shown that the presence of such cues decreases the size of place fields (Battaglia et al., 2004; Aikath et al., 2014; Zhang et al., 2014).

Compared to real-world free exploration, place coding in the floating track system does not persist for multiple days to the same degree, similar to VR systems. Another limitation of the floating track system compared to real-world free exploration is that it is constrained to relatively small environments – the longest track we used in this study was 1 metre in length, although we have designed environments that work within the current setup that allow tracks as long as 2 metres. Although not as long as the 18 metre tracks of Brun et al. (2008), this should be well within the ecological regime for mice. In the floating track, distal visual cues are absent. As it is the track that moves, not the mouse, any distal cues would present a confounding stimulus. When distal visual cues are minimal and proximal cues are prominent, it has been shown that the proximal stimuli significantly influence place cell activity (Young et al., 1994; Shapiro et al., 1997). In the floating track, proximal cues play an important role in influencing place cell activity. Place field locations are strongly tied to the proximal cues with more place cells mapping positions near the cues. Further, when the proximal cues are changed, the place fields remap. The floating track system has one distinct advantage over real-world behavioural tasks in allowing head-fixation, making it compatible with a wider range of optical imaging technologies and allowing for straightforward extension to intracellular recording.

This study focused on CA1 hippocampal place cell representations during exploration. The floating track system presented here offers advantages over traditional VR systems and free exploration real-world systems for investigating neural encoding in a variety of brain systems, and may well prove a useful tool for dissecting neural pathways underpinning cognitive functions based upon spatial representations, such as spatial working memory tasks (Lalonde, 2002; Dudchenko, 2004) and object location and recognition tasks (Vogel-Ciernia and Wood, 2014). Floating track behavioural tests may occupy a useful position between virtual reality paradigms, which allow head-fixation but are generally limited to simplistic unimodal sensory stimulation, and real-world behavioural tasks.

We found that approximately 88% of the CA1 cells we detected were active (fired at least once) in any given session (see Fig. 5C). If a cell was active in a given session, then it was around 90% likely to be active in any subsequent session, with no obvious temporal dependence in a timescale of several days. Approximately 29% of detected cells were place-sensitive (see section 3.2). One caveat concerning multiphoton calcium imaging in sparsely firing areas such as hippocampal region CA1 is that accurate identification of the boundaries of ROIs corresponding to single cells depends upon the cells firing at least once. In our data analysis here, we used the CaImAn software package (Giovannucci et al., 2019) to extract ROIs, together with extensive modifications to the data processing pipeline to improve registration of ROIs across sessions on our dataset, as well as incorporating additional neuropil decontamination. We have also used ABLE (Reynolds et al., 2017) in place of CaImAn in this pipeline with similar results. In both of these software tools, cells are registered only if they are active at some point in the set of sessions imaged. This suggests that we are likely to be under-estimating the total number of neurons in the field of view, and thus over-estimating the fraction of active and place-tuned cells. It should also be noted that the non-place-tuned cells we observed may include cells that might show place tuning in a freely moving (non head-fixed) scenario. We additionally note that comparison of results from multiphoton imaging with electrophysiological studies can be complicated by both inherent biases in cell selection, as described above, and by the different observation model due to calcium measurements, which has been suggested to lead to a lower fraction of responsive neurons, but sharper selectivity (Siegle et al., 2020).

The visual stimulus apparatus we used in this study consisted of phosphorescent tape formed into patterned structures on the walls of the floating track. This resulted in scotopic illumination conditions, which correspond to decreased spatial acuity (Umino et al., 2008). However, this is not a fundamental aspect of the platform, and instead photopic illumination could have been used, together with higher walls and a cone around the objective lens to exclude light from the microscope emission path.

A further caveat to our study is the difference in the age and gender profiles of the mice imaged in the circular track and open field tasks, with younger, female mice used in the latter task. While there is data suggesting no gender effect should be expected (Fritz et al., 2017), we might expect lower spatial selectivity and reduced place field stability (Yan et al., 2003) in the older mice we imaged in the circular track. We can, thus, view our results in the circular track as conservative with regard to the tuning and stability that might be expected in younger mice, while demonstrating the applicability of the platform to the study of neurodegenerative models which require the use of older mice.

The idea that coding in CA1 and other brain areas is dynamic – i.e. that the ensemble of cells encoding a given memory evolves over time – is now relatively well-established (Ziv et al., 2013; Rubin et al., 2015; Hainmueller and Bartos, 2018). Our results suggest that the time constant of this reconfiguration may depend upon environmental factors and the number and type of sensory cues being integrated, with head-fixed preparations including virtual-reality and floating track paradigms inducing faster reconfiguration, while memories formed during freely moving behaviour may induce cell assemblies that are stable for longer durations, as observed in Ziv et al. (2013). (However, noting the caveat above, it is possible that greater place field stability would have been observed in younger mice). Interestingly, similar changes in the specific ensemble of neurons encoding a sensory stimulus have been observed in the olfactory bulb (Kato et al., 2012), whereas motor cortex seems to show no such effect (Peters et al., 2014). This underscores the importance of further research on the specific parameters affecting the stability of neural representations employed in perception, memory and cognitive behaviour, with technology such as that described here now making practical the longitudinal experiments necessary to address this issue.

The relatively high reproducibility of stimulus conditions in the circular track task allowed us to make use of a trial-to-trial variability analysis method widely employed in visual neuroscience, which has however not, to our knowledge, previously been employed to study neural representations in spatial memory. This led to the interesting result that while place cells in CA1 fire at higher rates than non-place cells, their activity is in fact more reliable from trial to trial, given the firing rate – i.e. that while neural response variance increases with the mean level of activity, it does so with a lower exponent for place cells than for non-place cells, indicating comparative reliability. We speculate that, like spatial information (Cacucci et al., 2008), the response variability exponent may turn out to be a sensitive indicator of neurodegeneration under conditions of aberrant excitability (Busche et al., 2008).

As well as examining single cell response properties, we used a manifold learning approach to study the behaviour of the entire ensemble of recorded cells during spatial exploration. We used the Multidimensional Scaling (MDS) manifold learning approach, after observing that it performed better than Principal Components Analysis (PCA) in terms of variance accounted for, however we verified qualitatively that a similar picture was provided by both PCA and Isomap (see Supplementary Fig. 3), and expect our results to be robust to any sufficiently comprehensive manifold learning approach. The spatial position of the mouse in the 1D (circular track) task was visually apparent in the lowest dimensions of the neural manifold; while this was less immediately obvious (because of the non-cyclical nature of the exploratory trajectory) in the 2D task, a decoding analysis indicated that for the open field task also, spatial location could be decoded with some success from a relatively small number of manifold dimensions, although with much less fidelity than for the circular track. As might be expected given the more constrained nature of the task, the dimensionality of the neural manifold obtained in mice exploring the circular track was lower than that observed in mice exploring the open field. The use of manifold learning in conjunction with a floating real-world environment may provide a useful tool for uncovering fundamental principles governing encoding for memory, perception and cognition.

A significant advantage of the floating arena is its application to behavioural tasks which cannot otherwise be performed with a head-fixed animal. Examples include object-in-place tasks (wherein real-world objects can be introduced to arena locations and physically explored by the mice, contextual fear conditioning (applied via a foot-cuff electrode or similar), spatial goal-oriented behaviour (which we explored in the early stages of developing our task, confirming the over-representation of place fields around a fixed reward zone), and social-context discriminations (possible by having a second mouse in another fixed or floating mini-arena on the platform), and a variety of other tasks involving conjunctive representations with a spatial component.

In this paper we have validated a platform technology which, by combining multiphoton calcium imaging in head-fixed mice with floating track implementation of a behavioural task, allows robust and sensitive readout of memory encoding and retrieval performance from the activity of hippocampal neurons. As well as enabling advances in cognitive neuroscience, we expect this tool to be of great utility for the study of mouse models of neurodegenerative disorders, and a powerful tool for aiding the pre-clinical development of therapeutics for these diseases.

## AUTHOR CONTRIBUTIONS

MAG and SRS designed the experiments. MAG worked out the technical details of the experiments. MAG and JR performed the experiments with support from YL. MAG processed the experimental data and performed the analysis with support from CD, GPG, SP and SRS. MAG and SRS wrote the manuscript. MAG and SRS supervised the project. All authors discussed the results and commented on the manuscript.

## ACKNOWLEDGMENTS

We thank David Dupret, Richard Morris and Leo Khiroug for useful discussions, and Sihao Lu for his assistance with the Matlab adaptation of the FISSA toolbox.

## SUPPLEMENTAL DATA

The Supplementary Material for this article can be found in the attached Supplementary Material file.

## Supplementary Material

**Figure S1.**
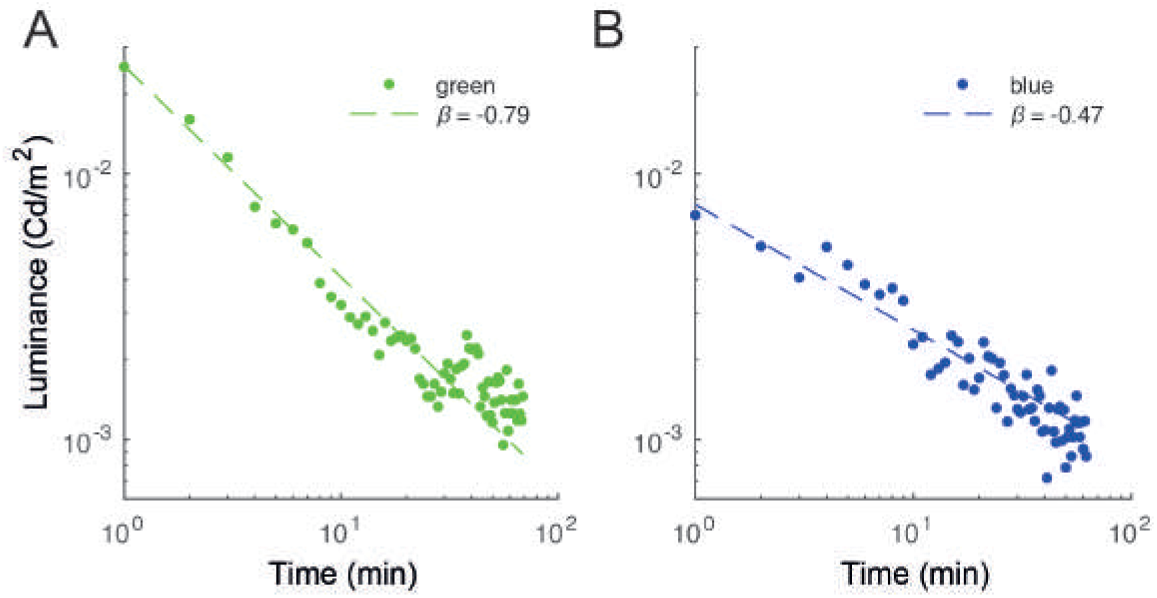
Luminance of phosphorescent tapes used as visual cues decays at a power law rate. Tapes were coloured (**A**) green (peak 520 nm) and (**B**) blue (peak 500 nm). Dashed lines denote curve fits of the form *y = αx^β^*. The luminance over an imaging period of an hour is well within the sensitivity range for mice, within the scotopic (low-light) range (Umino et al., 2008).

**Figure S2.**
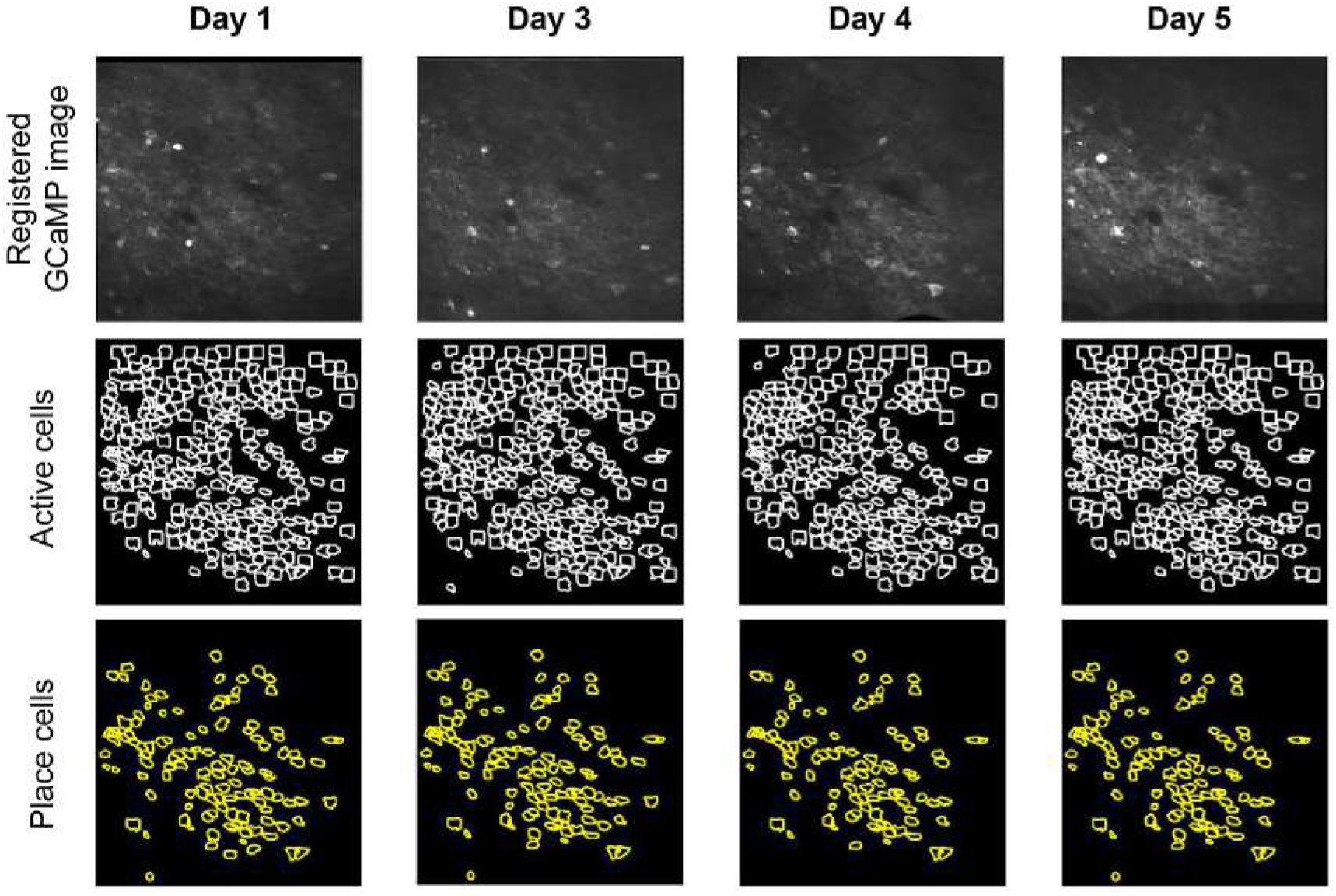
Image registration allows cells to be tracked across multiple imaging sessions. GCaMP images from different sessions were motion-corrected using the image from one session as template. The registered images (top row) were then temporally concatenated and the ROI segmentation algorithm was run on the concatenated video to produce a map of all the active cells. Active cells for each session (middle row) were identified based on the deconvolved neural activity for the session. The subset of active cells that were place-sensitive (bottom row) were then identified by place field analysis.

**Figure S3.**
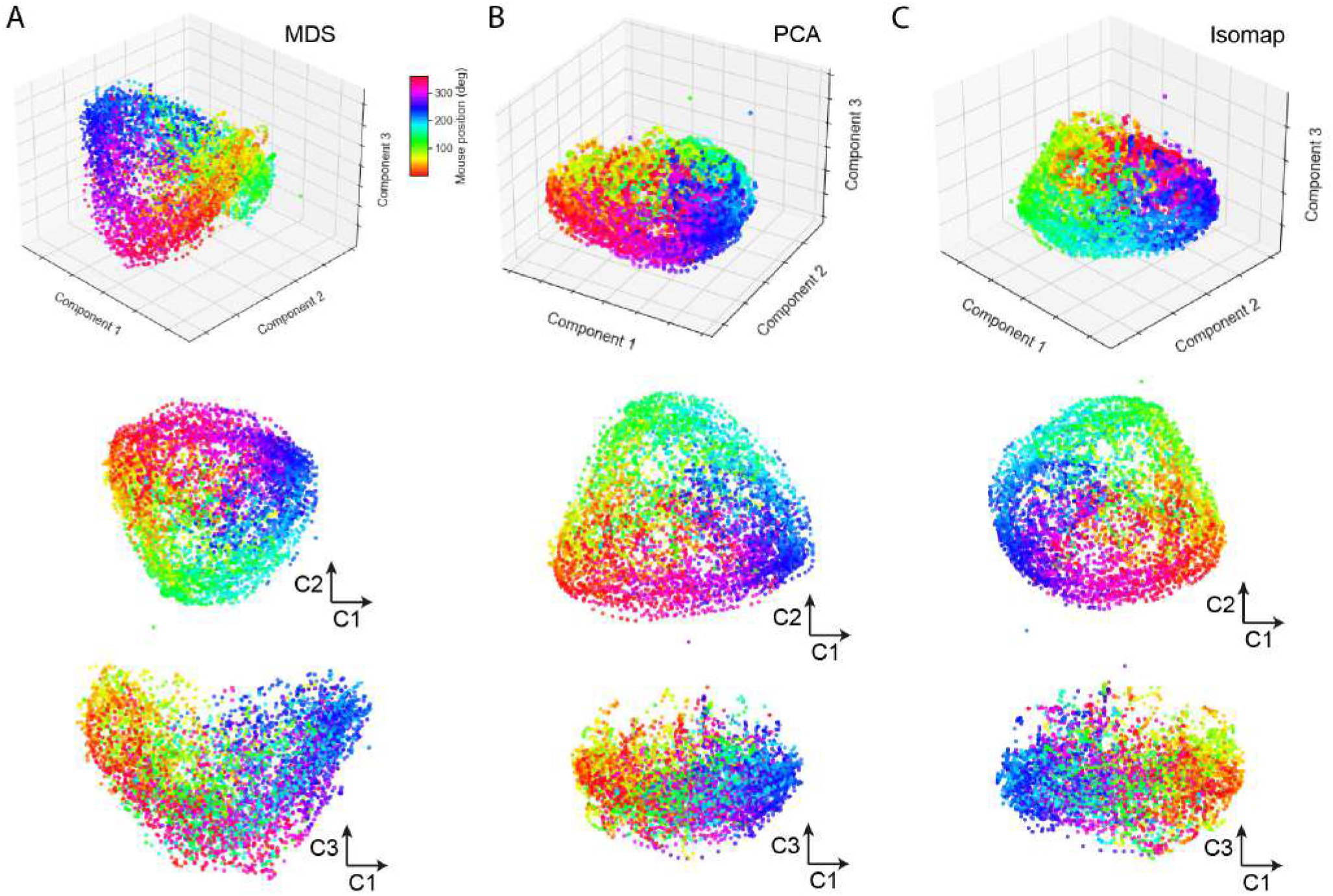
Neural manifolds extracted using (**A**) MDS, (**B**) PCA and (**C**) Isomap are very similar. Example shown here is the same circular track recording shown in the main manuscript. While MDS systematically captures at least as much of the variance as PCA (normally more) for the same number of dimensions, it also shows residual sructure related to mouse spatial position beyond the second dimension, which is not apparent in PCA or Isomap. Isomap provides a very similar depiction to MDS and PCA, although the ‘‘variance accounted for” quantification of manifold performance does not directly apply in this case. Our conclusion is that the precise technique used to visualise the manifold is not crucial.

